# Phenotypic screening for small molecules that lower PrP in cultured cells

**DOI:** 10.64898/2026.04.07.716919

**Authors:** Jeannine A Frei, Andrew G Reidenbach, Leo MH Xu, Raja Mohan Gopalakrishnan, Dominick Casalana, Daniel A Sprague, Mark-Anthony Bray, Amy Q. Wang, Vanessa Laversenne, Brian Erickson, Craig Braun, Samarjit Patnaik, Mckenzie Hall, Douglas Auld, Eric Vallabh Minikel, Sonia M Vallabh

## Abstract

PrP lowering is a validated therapeutic hypothesis in prion disease. To identify small molecules that reduce PrP levels, we performed phenotypic screening in cultured cells. To prioritize PrP specificity in our primary screen, we generated mouse N2a cells stably expressing GFP and used high content imaging analysis to select compounds that lowered PrP without affecting GFP signal or cell viability. Screening a curated library of 3,492 compounds with annotated mechanisms of action identified two small molecules, EYH (PubChem CID: 71678945) and LCZ (PubChem CID: 24970350), that selectively and dose-dependently lowered PrP. Proteomics on whole cell lysates identified PrP as the #1 or #2 most potently downregulated out of 8,722 proteins detected. Both compounds minimally affected *Prnp* mRNA, reduced expression of exogenously transfected PrP, and remained potent in non-dividing primary cells, consistent with a post-translational mechanism. Co-treatment with the proteasome inhibitor MG132 yielded accumulation of unglycosylated PrP, demonstrating proteasome clearance of PrP. However, both compounds showed limited or no activity in human cell lines, and failed to reduce PrP in vivo after 14 days of treatment. These findings highlight the challenges associated with mechanism-agnostic phenotypic screening for PrP-lowering compounds and support prioritizing compounds with known mechanisms of action.

## Introduction

PrP lowering is a validated therapeutic hypothesis in prion disease^1^. Prion disease is a rapidly progressive neurodegenerative disease of humans and other mammals caused by accumulation of misfolded conformers of the endogenous PrP protein, whose abundance dose-dependently drives prion replication and disease progression^2^. The relationship between PrP expression level and prion disease tempo has been demonstrated in genetic models^3,4^ and across multiple pharmacologic interventions, including antisense oligonucleotides (ASOs)^5,6^, a divalent siRNA^7^, a CRISPR-guided base editor^8^, and a zinc finger repressor^9^. Notably, an ASO reducing PrP recently completed a Phase 1/2 trial in symptomatic patients diagnosed with prion disease^10^. However, these platform approaches rely on either intrathecal administration, which yields broad but uneven CNS biodistribution, or on successful translation of novel viral vectors engineered to bind human receptors. In contrast, small molecules, which may diffuse through tissue without compartment-restricted or receptor-mediated delivery, represent an appealing additional therapeutic modality.

An unbiased RNA interference screen identified multiple genes regulating PrP expression, and some of them may represent druggable targets^11^. Several prior high-throughput screens attempted to identify small molecules that lower PrP^12–15^, yet none have demonstrated in vivo efficacy. Several limitations of these efforts are apparent. Pan-assay interference compounds (PAINS) can dominate the screening hits^16^, although curated compound libraries can mitigate this risk. In addition, PrP phenotypic screens in dividing cells readily identify non-specific compounds that broadly suppress protein expression. Cytotoxicity counter-screens may be insufficient to exclude such compounds, whereas orthogonal protein counter-screens and unbiased proteomics can assist with prioritization. Reproducibility also remains challenging, for instance, in one approach, astemizole the top hit in a pilot screen of the US Drug Collection^15^, but proved inactive when screened as a larger compound library in the main screen^17,18^. Increased technical replicates and/or parallel screens across multiple cell lines may help address such issues.

Incorporating the lessons learned from past studies, we designed a phenotypic screening campaign to identify PrP-lowering small molecules. We generated stable N2a cell lines expressing either cytosolic GFP or GFP-GPI, a GPI-anchored protein with a similar PrP biosynthesis pathway, thus enabling a built-in counter-screen for non-specific compounds across multiple cell lines. Using a high content imaging platform, we quantified PrP and GFP expression along with cell count to assess cytotoxicity while leveraging replicates to improve robustness. We then screened the curated “MoA box”, a library of 3,492 compounds with annotated mechanisms of action^19^.

Here we describe the design and execution of this screen and the identification and characterization of two relatively selective PrP-lowering compounds. Despite the careful design of our screen, neither compound demonstrated an acceptable therapeutic index in human cells, nor evidence of in vivo activity, and a precise exact molecular mechanism of action could not be established. These findings highlight the challenges of this approach but also offer support for continued investment in platform therapeutics with defined mechanisms of action and in vivo delivery efficacy.

## Results

### A high-content imaging phenotypic screen identifies three PrP-lowering compounds

Mouse N2a cells stably expressing cytosolic GFP or GFP-GPI were treated with test compounds in 1536-well plates at 1 µM or 10 µM in technical duplicates, yielding 8 wells per compound (Figure 1A). 50 nM PrP-targeting siRNA was used as positive control. In a primary screen of 3,492 compounds from the MoA box, technical replicate correlation coefficients per concentration were 0.795 for cytosolic GFP and 0.789 for membrane PrP (Figure S1A). We selected 359 compounds for re-testing, of which 337 of these confirmed activity and were advanced to 4-point dose-response. 44 compounds were subsequently advanced to 8-point dose-response scanning 6 nM to 20 µM (Figure 1B). The top 10 compounds with a 10-fold or greater therapeutic index between the IC_50_ at which PrP was lowered and the IC_50_ at which GFP signal and/or cell viability were reduced, were tested by Western blot in untransfected N2a cells using resupplied material for confirmatory testing when available (Figure 1B-C, S1B, supplementary table 1). Of the 10 hits tested by Western blot at 2 µM, 5 showed no discernible activity, while 2 (TCS 21311 and autophinib) showed marginal activity. Three compounds potently reduced PrP; EYH and LCZ uniformly diminished di-, mono-, and unglycosylated PrP, while Y-320 lowered PrP expression and impacted glycosylation, causing a downward shift in the molecular weight of the diglycosylated PrP band (Figure 1C). All three compounds dose-dependently lowered PrP, with substantial reduction at 5 µM and 10 µM (Figure 1D). 8-point dose-response curves showed EC_50_ values of 2 µM for EYH, 0.5 µM for LCZ, and 0.2 µM for Y-320 (Figure 1E). Immunofluorescence images from the original screen confirmed that all three compounds reduced PrP visually to nearly undetectable levels, similar to 50 nM PrP siRNA, without obvious impact on GFP-positive cells or on cell density (Figure 1F). The structures of the three compounds revealed no obvious pan-assay interference liabilities (Figure 1G). These three compounds were selected for further characterization.

**Figure 1.**
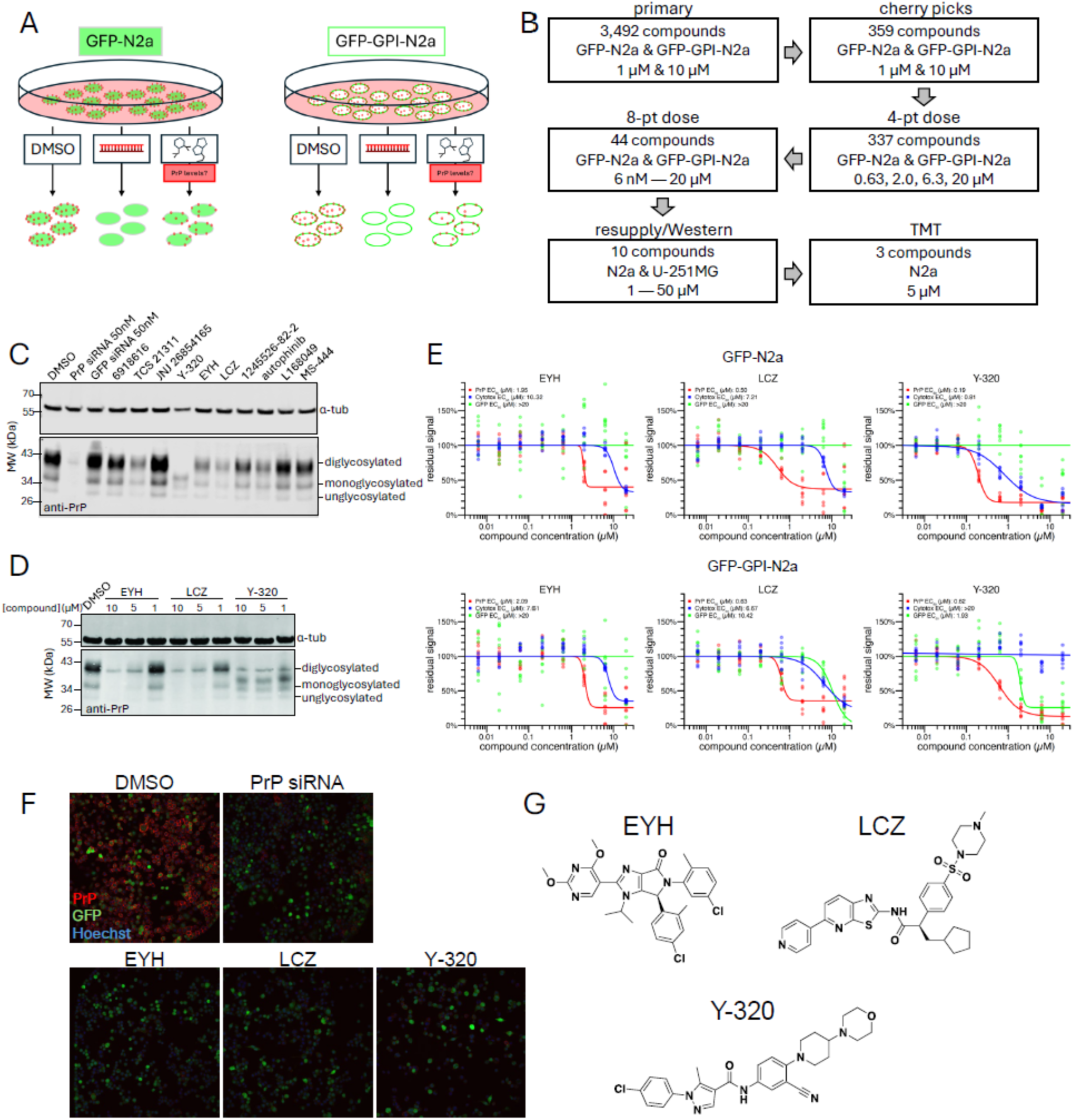
Discovery of PrP-lowering compounds. **(A)** Schematic overview of the phenotypic screening strategy for PrP-lowering small molecules. N2a cells stably expressing GFP or GFP-GPI were treated with DMSO, Prnp-targeting siRNA or test compounds. PrP expression was quantified by immunofluorescence to identify active compounds. **(B)** Compound triage flowchart. Primary screening hits were progressively narrowed through cherry pick selection, eight- and four-point-dose confirmation and Western blot validation. N2a cells treated with the three top candidates were analyzed by tandem mass tags (TMT) mass spectrometry. **(C)** Western blot of PrP expression in N2a cells treated with 2 µM of the ten compounds identified by the 8-pt dose response experiment. **(D)** Western blot of PrP expression in N2a cells treated with the three most potent PrP-lowering candidates EYH, LCZ and Y-320 at 1, 5 and 10 µM. (C, D) PrP detected with anti-PrP antibody POM2 (1:2000), anti-α-tubulin antibody (1:3000) served as loading control. **(E)** Eight-point dose response curves for the three top candidates in GFP-N2a and GFP-GPI-N2a cells, with PrP expression (red), cytotoxicity (blue) and GFP (green) as a function of compound concentration. **(F)** Representative immunofluorescence images of N2a cells stained for PrP (red), GFP (green) and Hoechst (blue) following treatment with DMSO, Prnp-targeting siRNA, EYH, LCZ and Y-320. All three compounds decreased PrP signal relative to DMSO-treated controls. **(G)** Chemical structures of the top candidate compounds.

### EYH and LCZ show high specificity for PrP

We performed tandem mass tag (TMT) proteomics on N2a cells treated with 5 µM compound or DMSO for 18 hours (Figure 2). After multiple testing correction (Bonferroni P < 0.05), 21 proteins (0.2% of those quantified) were significantly altered by EYH (Figure 2A), 51 (0.6%) by LCZ (Figure 2B), and 2099 (24%) by Y-320 (Figure 2C). Among Bonferroni-significant hits ranked by effect size (log2-fold change), PrP (gene symbol *Prnp*) was the most downregulated protein for EYH and the second most downregulated protein (after *Adam23*) for LCZ. In contrast, Y-320 reduced PrP by ∼41%, but this did not reach Bonferroni significance, with a *P*-value ranking 5,981st (31st percentile) across all proteins. These results indicate that Y-320 broadly modulates a large proportion of the proteome, whereas EYH and LCZ yielded greater specificity for PrP and were prioritized for further investigation.

**Figure 2.**
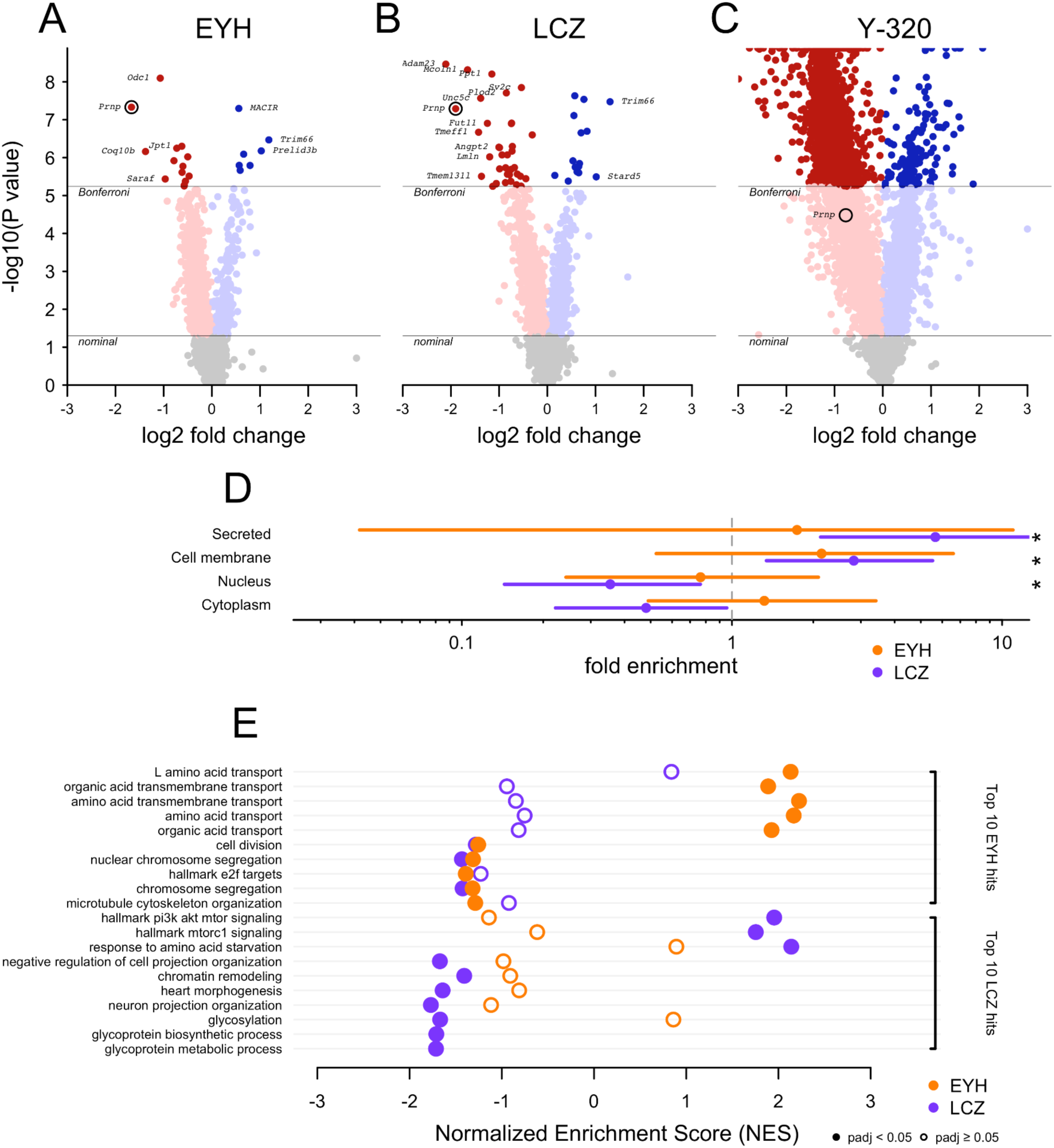
Proteomic selectivity of EYH and LCZ but not Y-320. N2a cells were treated with DMSO (N=6 technical replicates) or one of the three hit compounds (N=4 replicates each) at 5 µM for 18 hours. Cell pellets were subjected to unbiased tandem mass tag (TMT) proteomics to yield quantification of 32,721 peptides from 8,722 unique proteins (grouped by gene symbols). **(A-C)** Volcano plots of the log2-fold change (x-axis) versus -log10 P-value (Empirical Brown’s Method, see Methods). Lines indicate thresholds for nominal and Bonferroni-corrected significance for **(A)** EYH, **(B)** LCZ, and **(C)** Y-320. **(D)** Enrichment or depletion of UniProt-annotated subcellular localization among Bonferroni-significant hits. Fold enrichment for EYH and LCZ was calculated using contingency tables across all gene symbols for subcellular localization. Data points indicate mean fold enrichment, bars indicate 95% confidence intervals, and asterisks represent Bonferroni significance (P < 0.05) across eight Fisher tests. Note that enrichment here means that proteins with this localization are more likely to be hits; the analysis is not conditioned on the directionality of those hits (up versus down upon compound treatment) **(E)** Top ten enriched Gene Ontology (GO) terms for EYH and LCZ, with normalized enrichment score (NES) on the x-axis; filled symbols indicate adjusted P < 0.05 and open symbols indicate adjusted P ≥ 0.05.

Other than PrP, the only Bonferroni-significant hit shared by EYH and LCZ was upregulation of *Trim66* (examined further below). We examined the UniProt^20^-annotated subcellular localization of the Bonferroni-significant hits, regardless of whether those hits were up- or down-regulated. EYH hits were not enriched for any particular localization. In contrast, LCZ hits were significantly enriched for membrane and secreted proteins (19/51 Bonferroni-significant proteins), all (19/19) of which were downregulated (Figure 2D). The only GPI-anchored protein identified among Bonferroni-significant hits for either compound was *Nrn1* for LCZ. Gene ontology (GO) term analysis identified largely distinct pathways for the two compounds: the top ten EYH pathways were primarily associated with upregulation of amino acid transport while the top ten LCZ pathways were characterized by downregulation of glycosylation and glycoprotein metabolism (Figure 2E). None of these pathways were shared between the two compounds. The top EYH hits also showed downregulation of cell division and chromosome segregation pathways, some of which were also significant for LCZ, potentially reflecting downstream cytotoxicity.

MoA box annotations and literature searches did not identify an obvious mechanism by which EYH and LCZ might lower PrP. EYH (also known as compound 15a) is a close analog of the Phase 1 oncology drug candidate siremadlin^21^, discovered as a binder of MDM2 (PDB: 6GGN), that blocks MDM2-p53 interaction^22^. However, we did not detect p53 in our proteomics data, consistent with prior reports that N2a cells are p53-mutant and unresponsive to an MDM2-binding peptide^23^. Moreover, the reported EC_50_ of EYH for MDM2 binding (800 pM) is three orders of magnitude lower than its EC_50_ for PrP lowering. LCZ (also known as LCZ960) activates glucokinase (GCK)^24^, but was likewise not detected in our proteomics data and is not expected to be expressed at significant levels in brain or neuronal cells. Together, these observations suggest that PrP lowering by EYH and LCZ is unlikely to be mediated by their previously reported mechanisms of action.

### EYH and LCZ lower PrP through protein degradation

RT-qPCR of N2a cells treated with EYH and LCZ revealed a very slight but significant decrease in Prnp mRNA transcript (88% and 79% residual Prnp, respectively; Figure 3A). These changes are much smaller than the compounds’ effect sizes on PrP protein levels, thus suggesting a principal mechanism of action downstream of transcription and mRNA processing. To further support this notion, RK13 cells, endogenously lacking PrP, were transfected with a plasmid containing the PrP open reading frame (ORF) under the CMV promoter, thus creating a system lacking any endogenous upstream transcriptional or translational regulation. Indeed, both EYH and LCZ efficiently lowered PrP in this simplified system (Figure 3B), suggesting that the PrP promoter or regulatory translational elements are not required for their activity. We next turned to investigate the compound’s involvement in protein turnover. PrP’s reported half-life in mouse N2a cells is a few hours^25,26^, but approximately 5 days in vivo^27^. We turned to explore the compounds’ activity in primary neurons as we expected that they more closely mirror the in vivo turnover kinetics compared to mitotically active N2a cells. We hypothesized that the compounds should exhibit rapid, less than 24 h PrP-lowering activity only if they clear existing PrP protein. In contrast, if they blocked new PrP protein synthesis, clearance would be slower and changes in PrP levels only detectable after days. We found that primary cerebellar granule neurons grown for 14 days in vitro (DIV) and treated with compounds for 24 h exhibited reduced PrP expression (Figure 3C), consistent with a post-translational mechanism of action. We next sought to investigate the two major protein degradation pathways: the lysosomal pathway and the ubiquitin-proteasome system. Bafilomycin A, an inhibitor of lysosomal alkalinization, did not diminish the effect of either compound (Figure 3D), arguing against involvement of lysosomal degradation in clearance of PrP. The proteasome inhibitor MG132, however, caused accumulation of a low molecular weight band of unglycosylated PrP when co-treated with EYH or LCZ (Figure 3E), suggesting that proteasomal clearance plays a prominent role in the activity of these compounds. Accumulation of unglycosylated PrP upon co-treatment with MG132 was previously reported for cotransin, decatransin, and other broad spectrum inhibitors of the Sec61 translocon that prevent cotranslational translocation of PrP’s signal peptide into the ER^28,29^. This mechanism of action appears unlikely for EYH because its proteomic data showed no enrichment for secreted or membrane proteins likely to bear a signal peptide (Figure 2D). LCZ did show substantial enrichment for such proteins, though the GO term analysis highlighting glycoproteins in particular suggests a more specific activity than signal peptide translocation.

**Figure 3.**
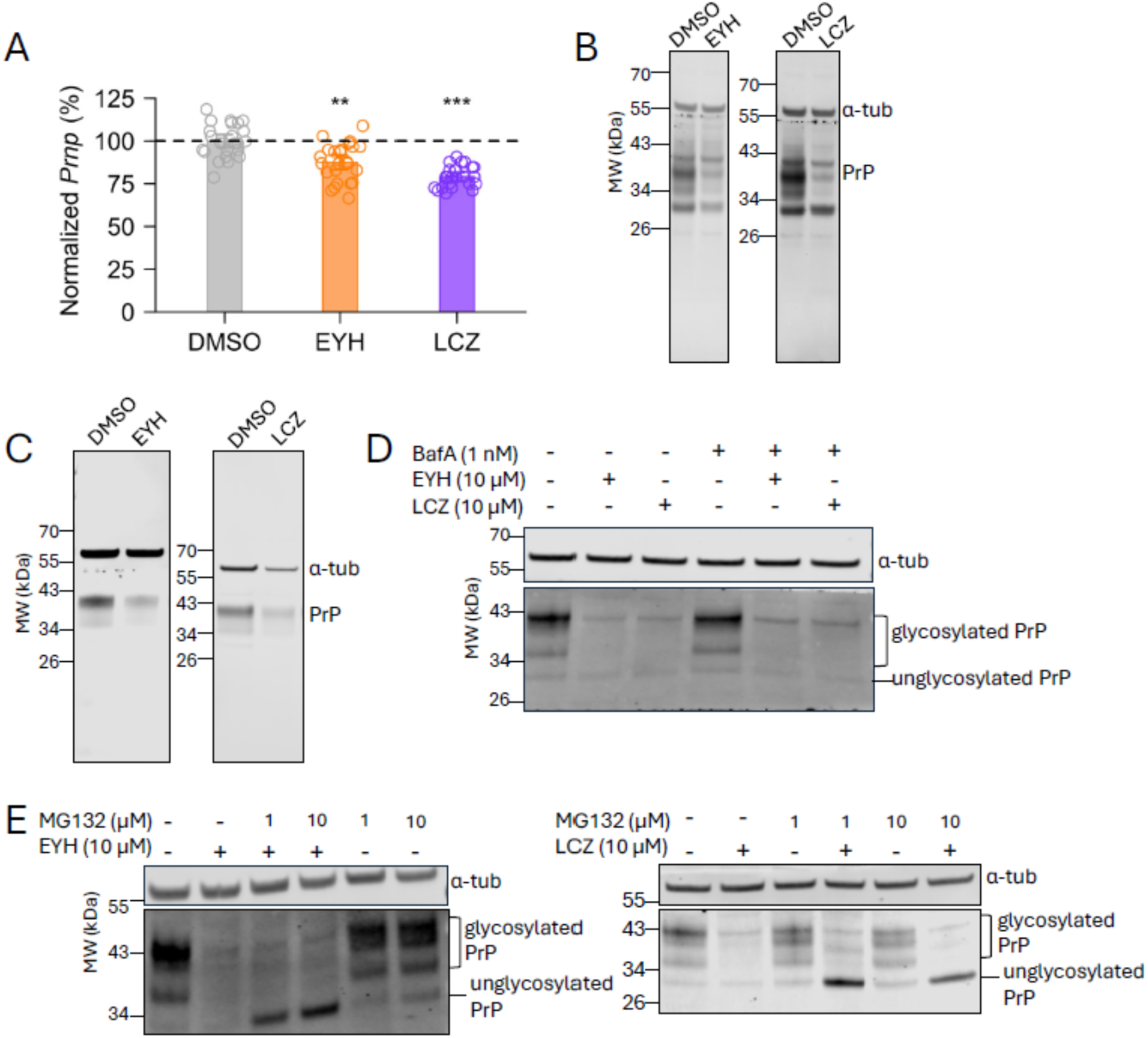
EYH and LCZ lower PrP post-transcriptionally through the ubiquitin-proteasome pathway. **(A)** Compounds exert a minimal effect on Prnp transcript. Prnp mRNA levels were analyzed by RT-qPCR. N2a cells were treated with 5 µM of EYH or LCZ, or DMSO only for 24 h. N=24, Kolmogorov-Smirnov test (** p < 0.01, ***p < 0.001). **(B)** ORF-transcribed PrP in natively non-PrP expressing RK-13 cells is responsive to EYH and LCZ. Cells were treated with 5 µM EYH or LCZ, or DMSO only for 24 h. Anti-PrP antibody 6D11 1:1000, anti-α-tubulin antibody 1:5000. **(C)** Compounds reduce PrP expression in non-dividing mouse cerebellar granule cells. Cells were cultured for 14 DIV and treated with 10 µM EYH or 6 µM LCZ, or DMSO only for 24 h. Anti PrP antibody 6D11 1:1000, anti-α-tubulin antibody 1:2000. **(D)** Inhibition of lysosomal pathway by BafA treatment does not rescue PrP lowering effect of compounds. N2a cells were co-treated with 1 nM BafA and 10 µM EYH or LCZ for 24hrs. Anti PrP antibody 6D11 1:1000, anti-α-tubulin antibody 1:2000 for EYH; anti PrP antibody 6D11 1:500, anti-α-tubulin antibody 1:3000 for LCZ. **(E)** Blocking the proteasomal pathway results in accumulation of cytosolic unglycosylated PrP. N2a cells were co-treated with 1 and 10 µM MG132 and 10 µM EYH or LCZ for 24 h. Anti-PrP antibody 6D11 1:500, anti-α-tubulin antibody 1:3000.

Trim66 upregulation was the only shared TMT hit among both compounds. We therefore asked whether Trim66 was mechanistically involved in PrP clearance (Figure S2). Both of two Trim66 peptides detected by TMT were upregulated to a similar extent in N2a cells treated with EYH or LCZ (Figure S2A) and both were found in all three UniProt-annotated isoforms of Trim66. We therefore focused on the canonical isoform 1. Both compounds increased Trim66 transcripts within 2 h of treatment, suggesting a transcriptional mechanism (Figure S2B). We were able to achieve both plasmid-driven overexpression (Figure S2C) and ∼40% siRNA knockdown (Figure S2D) of Trim66 in N2a cells. However, Trim66 overexpression did not impact PrP levels (Figure S2E), and neither overexpression nor knockdown of Trim66 altered the potency of EYH or LCZ (Figure S2F, G).

### Diminished activity of compounds in human cell lines

We envisioned that a CRISPR loss-of-function screen in human cells stably expressing Cas9, treated with EYH or LCZ, and flow-sorted for undiminished PrP expression would be an effective way to identify effector genes responsible for the activity of EYH and LCZ. Because human U251-MG glioblastoma cells express PrP and have been useful for screening siRNA sequences^7^, we tested whether EYH and LCZ were active in these cells. Both by flow cytometry (Figure 4A) and by Western blot (Figure 4B), we observed that each compound lowered PrP only at concentrations of 20 µM or above, an order of magnitude higher than the EC_50_ in N2a cells, with significantly affected cell viability at these concentrations (Figure 4B). This indicates little or no therapeutic index. Interestingly, EYH treatment at 50 µM not only resulted in clearance of glycosylated PrP, but also accumulation of unglycosylated PrP (Figure 4B), even in the absence of MG132, supporting a role in proteasomal clearance. We tested a panel of other human cell lines for compound responsiveness, all stably expressing Cas9 (except HEK293; Figure S3B) to potentially be used in a CRISPR screen. By flow cytometry, EYH showed no obvious activity in HEK293 cells at 10 µM (Figure 4C). At 50 µM EYH a small population showed reduced PrP expression, however cytotoxicity was pronounced. Among stably Cas9-expressing cell lines, EYH and LCZ exhibited some activity in A375 cells, albeit with lower potency compared to N2a cells, while activity appeared absent in HT29 cells and marginal in U87MG cells at the highest concentration tested (Figure 4D). Overall, no human cell line met all the criteria necessary to advance in the CRISPR screen (Figure S3A).

**Figure 4.**
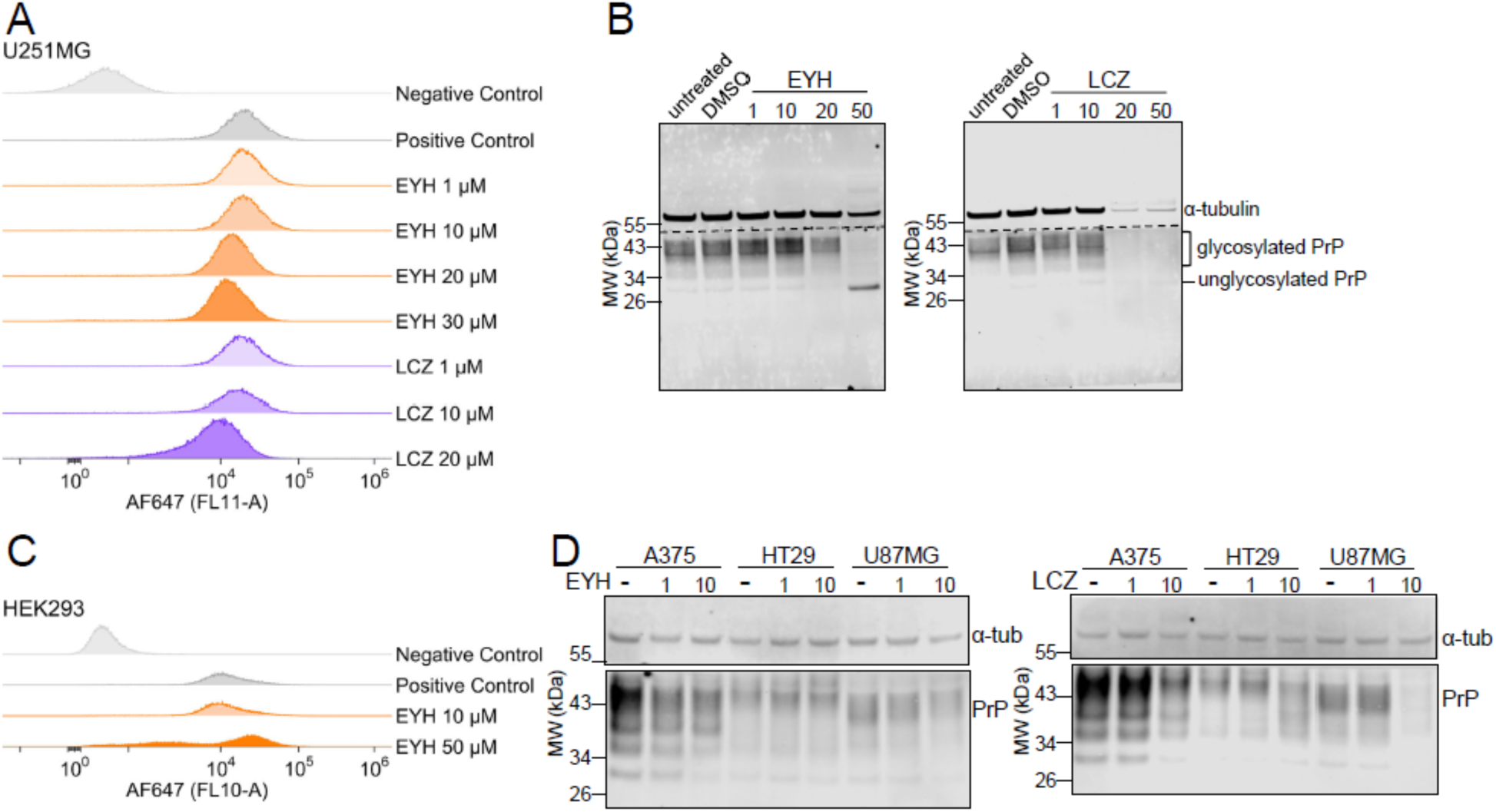
Compounds exhibit variably diminished potency across human cell lines. **A)** Flow cytometry histogram of PrP expression in U251MG-Cas9 cells treated with EYH (orange) and LCZ (purple). Neither compound decreases PrP expression significantly in U251-MG cells. **B)** Western Blot of PrP expression in U251MG-Cas9 cells treated with 1, 10, 20 and 50 µM EYH and LCZ, respectively. PrP signal was decreased at 20 µM and 50 µM EYH, while unglycosylated PrP (∼34 kDa band) increased at the latter concentration. LCZ didn’t exert a PrP-lowering effect. Cell toxicity was observed at 50 µM EYH and above 20 µM LCZ. Anti-PrP antibody POM2 1:2000, anti-α-tubulin antibody 1:2000. **C)** Flow cytometry histogram of EYH-treated HEK293 cells measuring PrP expression. No significant reduction in PrP expression was observed at any concentration. (A and C) PrP expression was analyzed using anti-PrP 6D11 Alexa Fluor 647 antibody. Negative control (light gray): DMSO-treated cells stained with an isotype control antibody (Alexa Fluor 647 mouse IgG2a, κ isotype); positive control (grey): DMSO-treated cells stained for PrP. **D)** Western blot of PrP expression in A375-Cas9, HT29-Cas9 and U87MG-Cas9 cell lines treated with 1 and 10 µM EYH and LCZ, respectively. Compounds exhibit variable and low potency across Cas9-expressing human cell lines. Anti-α-tubulin antibody 1:3000, anti-PrP POM2 antibody 1:2000.

### Compounds do not exhibit PrP-lowering activity in vivo

We sought to test whether EYH and LCZ lowered PrP in vivo. Wild-type mice (N=7-8 per group) were dosed by oral gavage with either compound as a 100 mg/kg suspension (see experimental procedures), every day for 14 days, and tissue was harvested 3-6 hours after the final dose. We quantified PrP in the brain as well as in colon, a reasonably high PrP-expressing tissue useful as a measure for peripheral target engagement^27^. Neither compound had any effect on brain PrP concentration (Figure 5A). In colon, variability was higher, and a numerical increase in PrP concentration in EYH-treated animals reached nominal significance; neither compound decreased PrP (Figure 5B). We also performed pharmacokinetic measurements on the same brain, colon and plasma (Figure 5C), as well spleen and quadriceps (Figure S4), while acknowledging that the timing of harvest hours after dosing may more nearly reflect peak, rather than trough, tissue concentrations. EYH concentration in the brain (1.7 µM) was near the compound’s EC_50_ (2.0 µM) in N2a cells, while colon concentrations were an order of magnitude higher (32.3 µM). LCZ concentration in the brain (1.75 µM) was above its EC_50_ in N2a cells (0.5 µM), while colon concentration was 2 orders of magnitude above the EC_50_ (35.2 µM). The mean concentrations of EYH and LCZ in plasma were 16.7 µM and 6.25 µM, respectively. EYH tissue to plasma concentration ratios were 0.1 and 1.9 for brain and colon, respectively. LCZ tissue to plasma concentration ratios were 0.3 and 5.6 for brain and colon, respectively.

**Figure 5.**
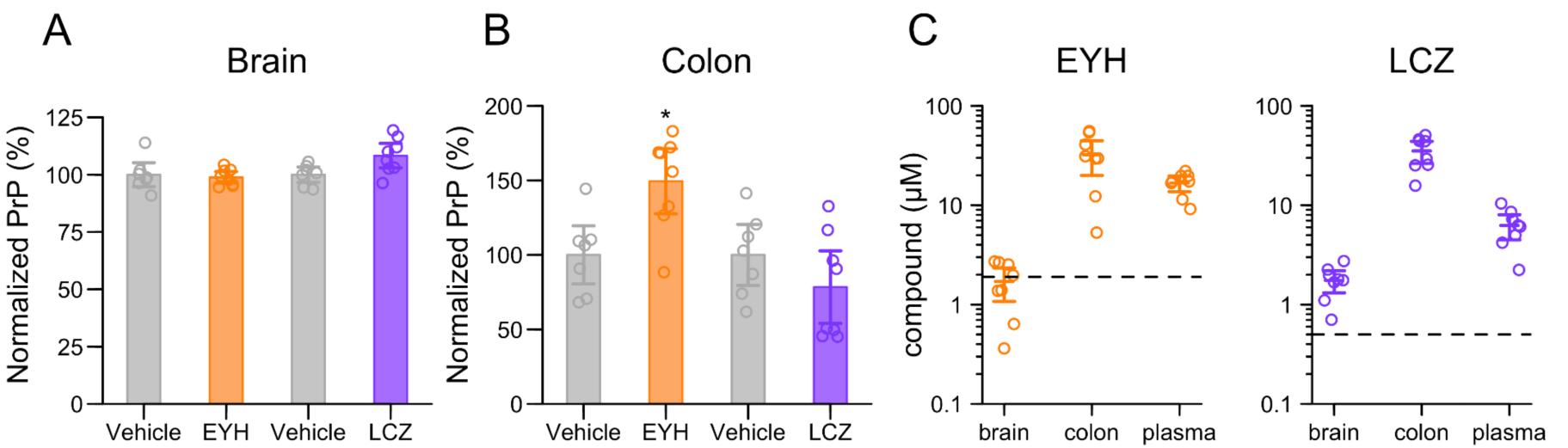
Lack of evidence for in vivo activity of EYH and LCZ. PrP expression in **(A)** brain and **(B)** colon of mice dosed with 100 mg/kg EYH (orange) and 100 mg/kg LCZ (purple). PrP concentration was measured by ELISA. Respective vehicle controls (grey) are shown to the left of each treatment group. No significant changes in PrP expression in brain was observed in mice treated with either compound. PrP was significantly increased in colon of mice treated with EYH, but not LCZ. Distributions of PrP between the control and treated groups were assessed using a Kolmogorov-Smirnov test (* p < 0.05). **(C)** Pharmacokinetics of EYH and LCZ in brain, colon and plasma. Dashed lines indicate EC_50_ of EYH (1.95 µM) and LCZ (0.5 µM) determined in N2a cells.

## Discussion

Despite incorporating lessons learned from prior phenotypic screens for PrP-lowering small molecules, we found that discovering biologically useful molecules in this fashion is very difficult. Our screening strategy was designed to select for reproducibility, selectivity, and interpretable, annotated mechanisms of action. We indeed discovered two small molecules that reproducibly and selectively lower PrP. Nevertheless, neither compound appears to be a tractable candidate for therapeutic development, and despite substantial follow-up studies, we were unable to determine a precise mechanism of action for either of them. Reflecting on this outcome reveals two fundamental limitations of the high-throughput screening paradigm: polypharmacology at screening concentrations, and model validity.

In virtually all high-throughput screens, including ours, compounds are screened at concentrations higher than those ultimately desired for a drug. In our case, we employed two concentrations — 1 µM and 10 µM — and utilized a carefully curated library of compounds, thus minimizing the impact of PAINS, which have plagued other high-throughput screens. Nonetheless, optimized compounds with well-characterized mechanisms of action, such as those in this screening deck, may exhibit picomolar or nanomolar potency for their characterized targets. When screened at thousands of times this concentration, these compounds may affect many other targets. This is both the reason why it is possible to find PrP-lowering compounds in a deck of only 3,492 small molecules, and the reason why such hits can prove difficult to advance therapeutically, or to assign a mechanism of action. Despite exhibiting quite high selectivity, the therapeutic index of our identified compounds in cell lines was only one order of magnitude, with concentrations higher than that leading to lowered GFP expression and affected cell viability. Meanwhile, the annotated mechanisms of action ultimately provided no indication towards the compound’s mechanism to lower PrP, given the irrelevance of the annotated targets in the cell lines used here and the vast difference in concentrations required for potency against previously annotated targets compared to PrP.

The ideal model for such a high-throughput phenotypic screen would be a PrP-expressing human neuronal cell line. In addition to us, previous studies report difficulties identifying human brain-derived cell lines that adequately express PrP protein for screening purposes^14,29,30^. N2a cells are murine, but are widely used in prion research for their abundant PrP expression and neuronal origin. We prioritized the expediency of executing a screen with N2a cells, and planned to validate hits in more clinically relevant models later. Unfortunately, the two compounds identified showed diminished potency in all human cell lines tested here, among many of them lacking a therapeutic index. Due to the lack of a compatible human cell line a loss-of-function screen using libraries of human CRISPR guide RNAs to identify effector genes responsible for the compound’s activity was abandoned. Meanwhile, although the compounds were active in mouse primary neuronal cells and had previously been reported in in vivo pharmacokinetic studies^22,24^, we were unable to identify in vivo target engagement, even with drug concentrations exceeding effective concentrations in some tissues, such as colon.

Previous PrP-lowering small molecule screens^13,14^ were reported at a time when technology to modulate the expression of one target protein in the brain was still in its infancy^31^. Small molecules, for all their challenges, appeared to have unique access to the human CNS. In the years since then, antisense oligonucleotides and siRNA have matured as drug modalities for the human CNS^32,33^ and have been shown to be effective in prion disease animal models^5–7^. Viral vectors designed to cross the human blood-brain barrier have been developed^34^, and cargoes such as transcriptional repressors, epigenetic editors, and base editors have been shown to lower PrP in mice^8,9,35^. A more rational, mechanism-based small molecule strategy of blocking passage through the Sec61 translocon in a signal peptide-specific manner is also reported to be in development^36^. Each of these approaches represent platform technologies with known mechanisms of action, and, thus, more predictable translational potential. Given the generally low success rate in drug development^37^ together with uncertainties about engineered viral vector translation to the human CNS and biodistribution of intrathecal oligonucleotides, investing efforts into mechanism-agnostic small molecule strategies initially seemed worthwhile. However, the roadblocks we encountered with EYH and LCZ underscore the profound challenges with even the most carefully designed mechanism-agnostic phenotypic screen. One could reasonably conclude, based on what we know today, that going forward the overwhelming weight of investment in drugs for prion disease should focus on rational platform approaches.

## Experimental procedures

### Cell lines and constructs for phenotypic screen

Plasmids encoding control proteins eGFP (MVSKGEELFTGVVPILVELDGDVNGHKFSVSGEGEGDATYGKLTLKFICTTGKLPVPWPTLVTTLTYGVQCFSRYPDHMKQHDFFKSAMPEGYVQERTIFFKDDGNYKTRAEVKFEGDTLVNRIELKGIDFKEDGNILGHKLEYNYNSHNVYIMADKQKNGIKVNFKIRHNIEDGSVQLADHYQQNTPIGDGPVLLPDNHYLSTQSALSKDPNEKRDHMVLLEFVTAAGITLGMDELYK) and GFP-GPI, with the signal sequence from *CD59* residues 1-25 and the GPI signal from human *CD59* residues 92-128 with attachment of the GPI anchor at residue N102. (MGIQGGSVLFGLLLVLAVFCHSGHSMVSKGEELFTGVVPILVELDGDVNGHKFSVSGEGEGDATYGKLTLKFICTTGKLPVPWPTLVTTLTYGVQCFSRYPDHMKQHDFFKSAMPEGYVQERTIFFKDDGNYKTRAEVKFEGDTLVNRIELKGIDFKEDGNILGHKLEYNYNSHNVYIMADKQKNGIKVNFKIRHNIEDGSVQLADHYQQNTPIGDGPVLLPDNHYLSTQSALSKDPNEKRDHMVLLEFVTAAGITLGMDELYKDLCNFNEQLENGGTSLSEKTVLLLVTPFLAAAWSLHP; Hiscox PMID: 12054528) were produced by Genscript with puromycin selection cassettes in the pSELECT-puro vector (Invivogen cat# psetp-mcs). We used Neuro-2a cells (N2a; ATCC CCL-131). After generation of puromycin kill curves, we determined that 3 µg/mL puromycin was necessary to kill all untransfected cells. N2a cells were transfected with Lipofectamine 3000 (Thermo L3000015) and 48 hours after transfection cells were selected with 3 µg/mL puromycin. Cells were grown in DMEM (high glucose, glutamine) supplemented with 10% FBS and 100 U/mL penicillin/streptomycin. Single clones were FACS sorted and allowed to grow into colonies. A single clonal line of the GFP and GFP-GPI cell lines were used for primary and secondary screening.

### Phenotypic screen

The primary screen was performed at 1 µM and 10 µM concentrations of compound in duplicates in 1536-well Aurora low profile plates (Aurora EBC241001A) with lids for automation (Greiner 691101). GFP and GFP-GPI cell lines were screened on separate days one week apart. 15 plates were used per screen. Plates 1 and 2 contained cells and DMSO, plates 3-14 contained compounds and plate 15 contained only DMSO. Columns 45-46 contained DMSO (n=64 wells per plate), column 47 contained 50 nM *Prnp* siRNA (n=32 wells per plate) and column 48 contained 50 nM *GFP* siRNA (n=32 wells per plate).

### Cell Culture

For compound screening studies cells were cultured in following media: N2a - DMEM, high glucose, GlutaMAX™ Supplement, pyruvate (Gibco 10569044); 10% FBS (Gibco 16000044); 100 U/mL Penicillin-Streptomycin (Life Technologies 15140163). U-251MG - Opti-MEM™ I Reduced Serum Medium, no phenol red (Gibco 11058021); 10% FBS (Gibco 16000044); 1X MEM Non-Essential Amino Acids Solution (Life Technologies 11140076); 1X GlutaMAX supplement (Life Technologies 35050079); 100 U/mL Penicillin-Streptomycin (Life Technologies 15140163). For mechanism of action studies N2a (ATCC, CCL-131), HEK293 (ATCC, CRL-1573) and RK-13 (ATCC, RK13) cells were cultured in DMEM/F-12 media (Gibco, 11320033) supplemented with 10% FBS (Gibco, 16000044) and 1% penicillin-streptomycin (Gibco, 15140163). Cas9 stably expressing cell lines (U251MG, A375, U87MG, HT29) were purchased from the Genomic Perturbation Platform at the Broad Institute. Cells were maintained in the following media supplemented with 10% FBS and 1% penicillin-streptomycin: U251MG and A375 cells in DMEM, U87MG cells in MEM, and HT29 cells in McCoy’s 5A media.

Primary cerebellar granule cell cultures were prepared from P6 to P7 C57BL/6N mice and cerebelli from three mice were pooled. In brief, cerebellum was dissected and meninges were removed. Tissue was enzymatically digested in papain (Worthington, LS003126) and cells were dissociated. Cells were plated at a density of 250,000/cm^2^ on plates coated with 20 µg/mL poly-D-lysine (Sigma, P6407). Cultures were maintained in serum-free Neurobasal A (Invitrogen, 10888022) media supplemented with 2 mM L-glutamine, 1% penicillin/streptomycin, 2% B27 supplement and 20 mM KCl for 14 days in vitro (DIV).

### Cell treatment

For the phenotypic screen, compounds (10 nL, Echo) and siRNA-lipofectamine mix (1 µL, manual pipetting) was preplated into a 1536-well plate and 1,500 cells/well (5 µL media/well) were plated on top of those reagents. Centrifuge the plate at 200 *g*, 1 min. Time from plating compounds to plating cells was about 2 h. Multidrop™ Combi Reagent Dispenser (Thermo) was primed with 70% EtOH, sterile water, and media before adding cells. After cell seeding, plates were incubated at 37 °C and 5% CO_2_ for 48 h. For mechanism of action studies, compounds were diluted in fresh media and added to the cells (0.2% DMSO end concentration). Immortalized cells were treated 24 h after plating, primary cells were treated at 14 days in vitro (DIV). All cells were harvested 24 h post-treatment. MG132 (Sigma, 474790) and Bafilomycin A1 (Cell Signalling Technology, 54645) were dissolved in sterile DMSO and diluted in media to 1 µM and 10 µM MG132 and 1 nM BafA, respectively. Cells were co-treated with 1 µM and 10 µM EYH and LCZ, respectively, and harvested 24 h post-treatment.

### Fixation

32% PFA (Electron Microscopy Sciences 15714-S) was diluted in DPBS (Thermo 14190144) to 8% PFA. 5 µL of 8% PFA was added to each well containing 5 µL of media (4% PFA final concentration) using a Combi on slow speed and incubated for 20 min at room temperature. Wells were washed three times with 8 µL DPBS using a Washer/Dispenser II (WDII; GNF Systems) and plates were stored overnight at 4 °C.

### Immunostaining

Wells were aspirated to 4 µL by WDII. A 4% BSA solution in DPBS was made and centrifuged at 7000 *g*, 5 min. 4 µL 4% BSA in DPBS was added on top of 4 µL of DPBS (Combi medium dispense speed) already in the plate and incubated 1 h at RT. WDII was used to remove 4 µL, leaving ∼4 µL in each well. 4 µL of primary antibody solution (anti-PrP 6D11 [Biolegend 808003] diluted 1:1000 in 2% BSA in DPBS) was centrifuged at 7000 g, 5 min and added (1:2000 final concentration, 1 µg/mL) and incubated overnight at 4 °C without agitation. Wells were washed three times in DPBS (8 µL) using the WDII, leaving ∼2-5 µL in each well. A master mix of Hoechst, CellMask and secondary antibody (2X concentrations in 4% BSA) was centrifuged at 7000x*g*, 5 min, and 4 µL of the supernatant was added to each well (Combi medium dispense, final concentrations anti-mouse Alexafluor 647, 1:2000 [Thermo A32728]; Hoechst 1:10,000 [Thermo H3570]; HCS CellMask™ Orange Stain [Thermo H32713] 1X, 1:5000). After incubating 60 min at RT protected from light, wells were washed three times with 8 µL DPBS using WDII. Approximately 10 µL of DPBS was added to the cells. Plates were sealed with foil cover and stored at 4 °C until ready to be imaged.

### Microscopy

Images were acquired with the Yokogawa CV8000 microscope (20X magnification). Two fields of view were taken from each well. Microscope exposure times were 500 ms PrP, 50 ms Hoechst, 50 ms GFP, 25 ms CellMask, all at 100% laser power.

### Image analysis

Both the GFP and GFP-GPI images were analyzed using CellProfiler^38^ version 2.2. Briefly, the nuclei were identified from the Hoechst image channel, using Otsu thresholding. Since the N2a neurons are tightly packed and proper identification of the boundary is important for membrane localization, the cell boundary was identified from the CellMask image channel by first smoothed slightly with a median filter to reduce noise. This image was edge enhanced in order to highlight the cell edges, and then subtracted from the original CellMask image; the resultant image had enhanced boundaries between cells to optimize detection. Using the nuclei as seeds, the cell borders were determined via Otsu thresholding followed by watersheding, and the cytoplasm was identified as the region between the nuclei and cell border. Misidentified objects were filtered based on their morphology. Subsequently, the cell membrane was identified as a 3-pixel region interior to the cell border. The raw PrP and GFP images were intensity-corrected by removing the background intensity using top-hat filtering. A suite of intensity-based readouts^39^ was collected from both the raw and corrected PrP and GFP images using the membrane and cell regions. We found that median intensity readouts using the background-corrected images performed better than the other image features.

### Hit calling

Treatments which reduced the cell count to <50% of the median DMSO cell count were excluded. In order to approach hit calling in an unbiased manner for the GFP and GFP-GPI cell lines, two methods were employed:

1. Well-level hits were called as those treatments which both decreased the cell membrane PrP WT levels (i.e., < 5th percentile of DMSO PrP intensity) and maintained membrane GFP WT levels (i.e., > 5th percentile of DMSO GFP intensity).
2. To leverage the siRNA knockdown (KD) treatments, a random forest classifier was created using the PrP KD and GFP KD single-cell phenotypes to create four classes for the training set, i.e., PrP+/GFP+, PrP-/GFP-, PrP-/GFP+ and PrP-/GFP-. Each well was then scored for the number of PrP-/GFP+ cells normalized by total cell count, and then Z-scored against the DMSO well distribution. Finally, single-cell hits were called as those treatments which increased the PrP-/GFP+ count ratio (i.e., > 99th percentile of DMSO count ratios).

Treatments that satisfied either hit-calling method were taken as primary hits and re-screened for confirmation.

### Four-point dose response

336 compounds were tested at 0.63, 2.0, 6.3, and 20 µM in quadruplicate.

### Eight-point dose response

44 compounds were taken into eight-point dose experiments at 6.3 nM, 20 nM, 63 nM, 200 nM, 630 nM, 2 µM, 6.3 µM and 20 µM in quadruplicate. Twelve of the 44 compounds were re-screened from the primary deck and were not present in the four-point dose experiment. These compounds had MoAs targeting OGT, GCK, GSK3B, ELAVL1, and MDM2, targets that appeared to be enriched in the positive hits. We also added two compounds whose MoA was against these targets but did not validate in the four-point dose experiment. The ten compounds that were effective in 8pt dose were taken into immunoblotting experiments.

### Compound resupply

Compounds EYH and LCZ are not commercially available and were provided by NIBR. 6918616 (23458), TCS 21311 (30063), JNJ-26854165 (16259), and Autophinib (27320) were purchased from Cayman Chemical; CAS 1245526-82-2 was purchased from Xcess Biosciences, Inc. (M10731-2); L-168049 was purchased from Tocris (2311); Y-320 was purchased from Selleckchem (S7516); MS-444 [AR6] was purchased from MedChemExpress (HY-100685).

### TMT

Tandem mass tag (TMT) proteomics data from 18 h compound treatments were analyzed for differential protein expression. Raw p-values from Empirical Brown’s Method^40^ (combining multiple peptide-level tests per gene) were adjusted for multiple testing using the false discovery rate (FDR) correction. Proteins were ranked by the metric |log2(fold-change)| × -log10(FDR-adjusted p-value), which prioritizes proteins with both large effect sizes and high statistical significance. Gene set enrichment analysis (GSEA) was performed using the fgsea R package^41^. Gene sets were obtained from the Molecular Signatures Database (MSigDB) v2024.1, including Hallmark pathways, Gene Ontology Biological Processes (GO:BP), and KEGG pathways. Pathways with 15-500 genes were tested. GSEA uses a signed ranking metric (preserving fold-change direction) to identify pathways enriched in upregulated or downregulated proteins. Pathway enrichment was considered significant at FDR < 0.05.

### siRNA and DNA transfection

Accell non-targeting control pool siRNA (Dharmacon, D-001910-10-05) and SMARTPool *Trim66* siRNA (Dharmacon, E-054730-00-0005) were reconstituted to 10 µM in 1x siRNA buffer (Dharmacon, B-002000-UB-100) according to manufacturer’s instructions. N2a cells seeded at a density of 1.5*10E4 were transfected with siRNA using Lipofectamine RNAiMAX transfection reagent (ThermoFisher, 13778030) according to the manufacturer’s protocol. 24, 48 or 72 h post-transfection, media was changed and cells were treated with 10 µM EYH or 10 µM LCZ for 24 h. *Trim66* open reading frame plasmid was generated by insertion of *Trim66* transcript variant 1 (accession no. NM_001170912) into pcDNA3.1(+) plasmid with a CMV promoter (GenScript) and N2a cells were transfected using Lipofectamine 3000 (ThermoFisher, L3000001) according to the manufacturer’s protocol. The PrP ORF (Genscript) was clone OMu23232 (accession NM_011170.3) in the pcDNA3.1-C-(k)DYK plasmid with a CMV promoter. Cells were harvested 24 h post-transfection.

### Immunoblots

For compound screening experiments, 1-2*10E5 cells/well N2a or U-251 MG cells were seeded in a 24-well plate. Cells were usually 50-80% confluent at time of lysis. Cells were washed once with 2 mL PBS and stored at −80 °C. Cells were lysed in 50-75 µL of RIPA lysis buffer with protease inhibitors per well (4 °C). For mechanism of action studies, N2a cells were seeded on 24 well plates at a density of 1.5*10E5. Human Cas9-expressing cell lines were plated on 12 well plates at following densities: 3.5-4.0*10E5 cells/well (U251MG), 2 wells pooled,: 1*10E5 cells/well (A375, 2 replicates pooled), 1.5*10E5 cells/well (HT29), 2.5*10E5 cells/well (U87MG). Cells were washed in ice-cold 1x PBS and lysed in ice-cold RIPA buffer supplemented with protease and phosphatase inhibitors (ThermoFisher, 78442), sonicated, and centrifuged at 4°C at 12,000 *g* for 5 minutes. Protein was quantified using the DC protein assay kit (BioRad 5000112). Lysates were mixed with 4x LDS (Thermo NP0007) supplemented with 50 mM TCEP (Thermo 77720). Protein-LDS samples were heated 5 min at 95 °C. Five micrograms of protein sample per lane were loaded on 4-12% NuPAGE gel (Thermo NP0323BOX, NP0321BOX) and run in 1X MES buffer at 40 min at 180V. Transfer to PVDF was performed on an iBlot2 at 20V, 7 min. The membrane was blocked with LICOR TBS blocking buffer (LICOR 927-60001) for 1 hr at RT followed by incubation of primary antibodies in LICOR TBS blocking buffer with 0.2%Tween overnight at 4°C. Primary antibodies were diluted to the following concentrations unless otherwise stated: 1:500-1:1,000 anti-PrP 6D11 (Biolegend, 808001); 1:2000-1;5000 anti-α-tubulin (Thermo, A11126), 1:3000 anti-LC3B (Novus, NB100-2220), 1:2000 anti-PrP POM2 (Millipore, MABN2298). Membranes were incubated with secondary antibody (IRDye 800CW Goat anti-Mouse IgG, LICOR, 926-32210; IRDye 800CW Goat anti-Rabbit IgG, LICOR, 926-32211) diluted in LICOR TBS blocking buffer (LICOR, 92760001), 0.2% Tween at a 1:10,000 dilution for 1 h at room temperature. Membranes were imaged on a LICOR Odyssey CLx Infrared Imaging System at 800 nm and 700 nm. Uncropped western blots are available in the online git repository of this manuscript.

### RT-qPCR

For RT-qPCR N2a cells were seeded at a density of 1*10E5 in triplicates on a 96 well plate. Cells were lysed and prepared for qPCR with the Cells-to-CT 1-Step Taqman Kit (Invitrogen, A25602) according to manufacturer protocol. TaqMan probes were used against mouse *Tbp* (Invitrogen, Mm00446971_m1), *Prnp* (Invitrogen, Mm00448389_m1), and *Trim66* (Invitrogen, Mm00725346_m1).

Each biological replicate was run in duplicate, normalized against the housekeeping gene *Tbp* and untreated control group (ΔΔCt). Samples were run in 384-well plates (Applied Biosystems, 430849) on a QuantStudio 7 Flex system (Applied Biosystems) with the following conditions: Reverse transcription (RT) 50 °C, 5 min; RT inactivation/initial denaturation 95 °C, 20 sec; Amplification 95 °C, 3 sec, 60 °C, 30 sec, 40 cycles.

### Flow cytometry

Cells were plated on 6 well plates at the following densities: U251MG: 1*10E6 cells/well, HEK: 5*10E5 cells/well, two wells pooled, U87MG: 7*10E5 cells/well, A375: 5*10E5 cells/well, HT29: 7*10E5 cells/well. Cells treated with EYH and LCZ were washed in PBS, gently detached and resuspended in Alexa Fluor 647 anti-CD230 (prion) antibody (clone 6D11, Biolegend, 808007) in flow buffer (1xPBS, 5% FBS). Primary antibody was incubated for 30 mins on ice. Cells expressing Cas9 were stained with Alexa Fluor 647 anti-Cas9 antibody (Cell Signaling Technology, 48796). In brief, cells were washed in PBS, trypsinized and resuspended in media. 1*10e6 cells were counted and resuspended in 50 µL chilled PBS. Cells were fixed and permeabilized using the Abcam Cell Fixation & Permeabilization Kit (abcam, ab185917). All cells except HEK329 were analyzed on a Cytoflex S flow cytometer (Beckman Coultier). HEK293 cells were analyzed on a Sony MA900 cell sorter (Sony). Nucleated cells were distinguished from debris and dead cells based on forward scatter-A/side scatter-A. Single cell population was distinguished from doublets and clumped cells based on forward scatter-A/forward scatter-H. Alexa 647 signal was detected by a 660 nm laser and 660/20 bandpass filters (Cytoflex S) or 638 nm laser and 665/30 bandpass filters (Sony MA900). Flow cytometry data was plotted in R version 4.4.3 using version 1.34.0 of the ggCyto package^42^.

### Mice

All in vivo experiments were approved by the Institutional Animal Care and Use Committee of the Broad Institute (Protocol #0162-05-17-2) and were performed in accordance with the National Institutes of Health *Guide for the Care and Use of Laboratory Animals*. Experiments in this study used C57BL/6N mice obtained from Charles River Laboratories. Mice were 13 weeks old at the time of administration of the first dose. Equal number of male and female animals were included in the study.

### Oral gavage

Our dosing approaches were guided by published in vivo work on these compounds. EYH (also known as compound 15a) was previously dosed orally at up to 5 mg/kg as a suspension in water with 0.5% carboxymethylcellulose and 0.5% Tween-80^22^. We formulated it in the same vehicle for 100 mg/kg dosing. At this concentration we did observe settling out of solids, and made an effort to repeatedly invert the mixture in order to ensure uniform dosing. LCZ (also known as LCZ960) was previously dosed orally at up to 300 mg/kg in plain water^24^. We formulated it for 100 mg/kg dosing in plain water, but despite the prior report, likewise found it incompletely soluble and made an effort to invert the mixture for uniform dosing. Mice were dosed by oral gavage with EYH (100 mg/kg, N=8), vehicle (DI water, 0.5% carboxymethylcellulose, 0.5% Tween-80, N=7), LCZ (100 mg/kg, N=8), or vehicle (DI water, N=7) for 14 days daily. 3-6 h after the last dose, blood was collected by cardiac puncture and brain, colon, quadricep and spleen were harvested and frozen on dry ice. All samples were stored at −80 °C until further processing.

### Homogenization

One brain hemisphere per animal was homogenized at 10% wt/vol and colon was homogenized at 10% w/v in cold 0.2% CHAPS in 1X PBS supplemented with one tablet protease inhibitor (Roche cOmplete 4693159001, Millipore Sigma, USA) per 10 mL in 7 mL (brain) and 2 mL (colon) tubes pre-loaded with zirconium oxide beads (Precellys, Bertin, USA) using 3 × 40 s pulses on a Bertin MiniLysis Homogenizer (Bertin, USA). Homogenates were aliquoted and stored at −80 °C until further analysis.

### ELISA

PrP concentration in the brain was measured using a PrP ELISA that has been described previously^43^. In brief, 96 well plates were coated with 2 µg/mL EP1802Y antibody (ab52604, Abcam, USA). After blocking in assay buffer (0.05% Tween-20, 5% BSA, 1X PBS), samples (10% w/v) were diluted 1:20 and incubated at RT for one hour. Biotinylated 8H4 antibody (ab61409, Abcam, USA) was added for one hour. Samples were incubated with streptavidin-HRP (Pierce High Sensitivity, 21130, Thermo Fisher Scientific, USA) for 30 minutes followed by addition of TMB (7004P4, Cell Signaling Technology, USA). The reaction was stopped by adding stop solution (Cat#) and the plate was read on a SpectraMax M5 platereader (Molecular Devices) at 450 nm and 630 nm. To quantify concentration of PrP in the colon, the same protocol as described above was used except that plates were coated with 0.25 µg/mL 6D11 antibody (Biolegend, 808001) instead of EP1802Y antibody. Colon samples were diluted 1:40 (10% w/v). Recombinant mouse PrP (MoPrP23-231) prepared as previously described^44^ was used to generate a standard curve. The concentration of residual PrP in each treatment tissue was divided by the mean concentration of residual PrP in control tissues to calculate knockdown of PrP protein.

### Bioanalysis and concentrations of EYH and LCZ in mouse plasma, brain, spleen, colon and quadricep

Drug concentrations in plasma and tissues were determined using an UPLC-MS/MS method developed by and implemented at NCATS. Ultra-performance liquid chromatography-tandem mass spectrometry (UPLC-MS/MS) method was developed to determine EYH and LCZ concentrations in mouse plasma, brain, spleen, colon and quadricep samples. Mass spectrometric analysis was performed on a Waters Xevo TQ-Absolute triple quadrupole instrument using electrospray ionization in positive mode with the selected reaction monitoring. The source settings for temperature, ion spray voltage and cone voltage were 600°C, 1.0 kV and 25 V, respectively. The MRM transitions were 591.2 →229, 552.2 →369.2 and 271 →155 for LCZ, EYH and tolbutamide (internal standard), respectively.

The separation of LCZ and EYH from endogenous components was performed on an Acquity BEH C_18_ column (50 x 2.1 mm, 1.7 µ) using a Waters Acquity UPLC system. The mobile phase A was 0.1% formic acid in water and B was 0.1% formic acid in acetonitrile. A gradient was run as follows: 0-0.2 min 10% B; 1.2 min 95% B; 2.0 min 95% B; 2.1 min 10% B at a flow rate of 0.6 mL/min.

Brain, spleen, colon and quadricep tissues were collected, flash frozen in liquid nitrogen and transferred to 48-well plates. The weight of each tissue sample was recorded. All plasma and tissue samples were stored at −80 °C until analysis. The calibration standards and quality control samples were prepared in the drug-free mouse plasma and tissue homogenates. Aliquots of 10 µL samples were mixed with 200 µL internal standard in acetonitrile to precipitate proteins in a 96-well plate. 1.0 µL supernatant was injected for the UPLC-MS/MS analysis. MassLynx and TargetLynx were used for data collection and processing (Waters Corp., Milford, MA).

### Statistics and data availability

Analyses used custom scripts written in R. All statistical tests were two-sided, and P values less than 0.05 were considered nominally significant. Data and source code will be made available at github.com/vallabhminikel/eyh_lcz

## Supporting information

This article contains supporting information.

## Acknowledgments

This work was funded by the National Institutes of Health (R01 NS131663).

## CrEDIT Statement

Conceptualization: DA, EVM, SMV; Data curation: AGR, MB; Formal analysis: JAF, AGR, LMHX, DAS, MB, EVM; Funding acquisition: SMV; Investigation: JAF, AGR, LMHX, RMG, MB, AQW, VL, BE, CB, SP, MH, DA; Methodology: JAF, AGR; Project administration: JAF, SMV; Supervision: SP, MH, DA, EVM, SMV; Validation: Visualization: JAF, AGR, LMHX, DAS, EVM; Writing – original draft: JAF, AGR, EVM, SMV; Writing – review & editing: JAF, AGR, LMHX, RMG, DAS, MB, AQW, VL, BE, CB, SP, MH, DA, EVM, SMV.

