## Supporting Information for "Phenotypic screening for small molecules that lower PrP in cultured cells"

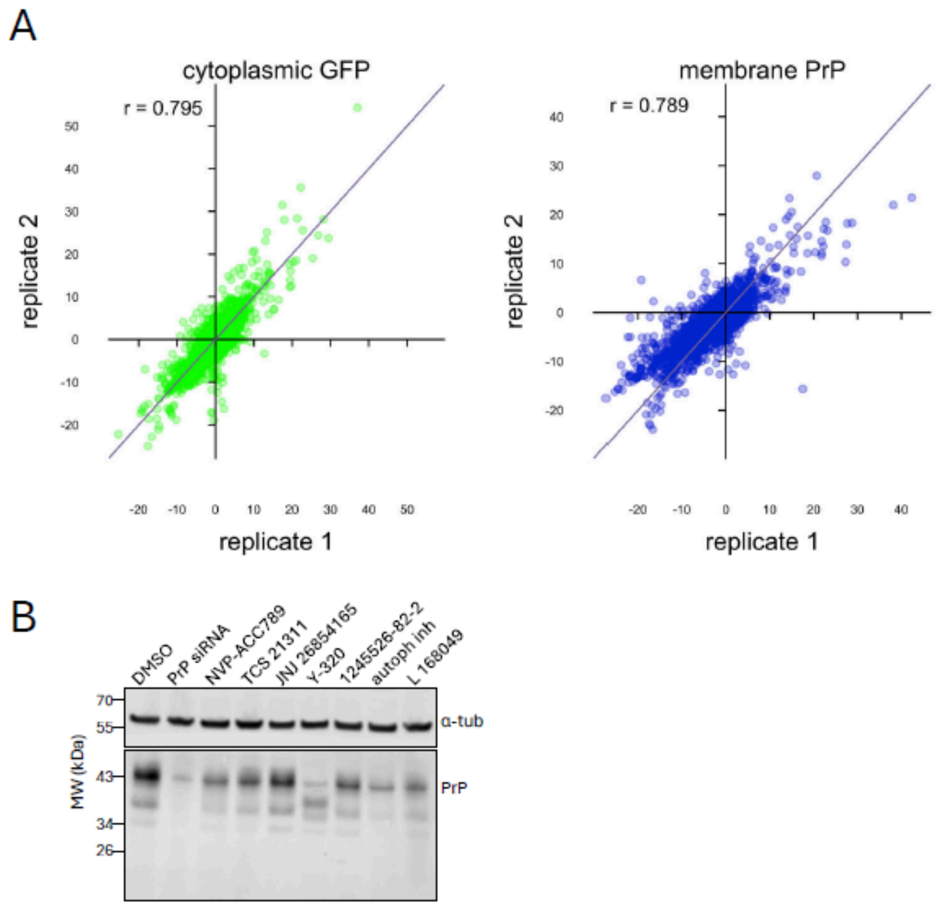


**Figure S1.** Development of the screen and validation of hits. **(A)** Reproducibility. **(B)** Western blot of PrP expression in N2a cells treated with 6 μM of commercially resupplied compounds identified in the 8-point dose response experiment.

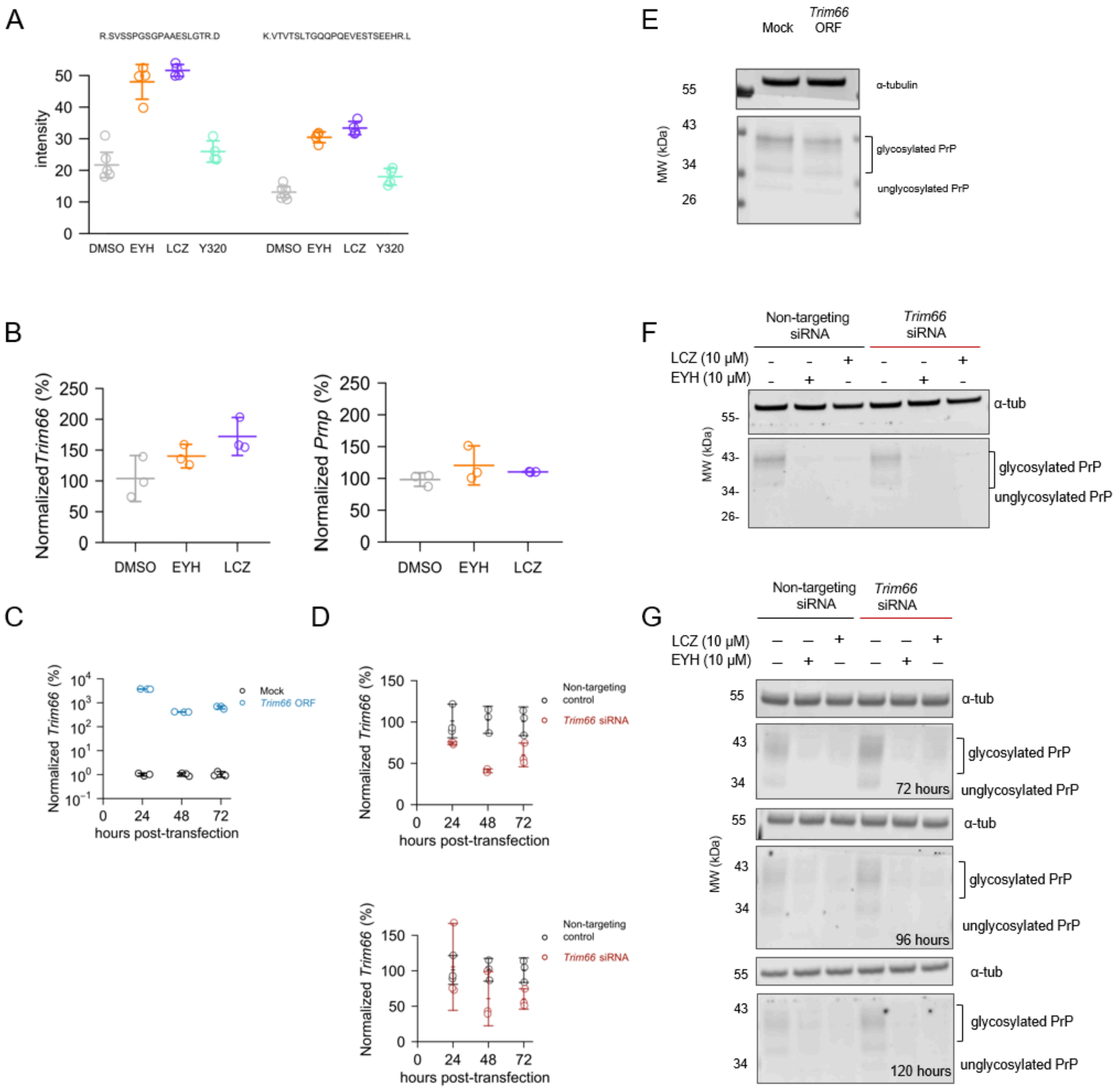


**Figure S2.** Trim66 has no obvious role in modulating PrP abundance. **(A)** Tandem mass tag mass spectrometry 18 hours post-treatment in N2a cells reveals upregulation of two peptide fragments found in all three isoforms of Trim66 after 5 µM of EYH and LCZ, but not Y320 treatment. **(B)** *Trim66* transcript levels trend upwards after 2 h of 10 µM treatment of EYH and LCZ, but *Prnp* levels are not affected. **(C)** Time course of *Trim66* isoform 1 open reading frame transfection into untreated N2a cells. **(D)** Time course of *Trim66* knockdown with 10 nM siRNA in N2a cells. **(E)** Overexpression of *Trim66* isoform 1 open reading frame in N2a does not affect PrP levels. N2a cells were harvested 24 hours post-transfection. **(F)** 24-hour pretreatment of 10 nM *Trim66* siRNA does not ablate the PrP-lowering effects of EYH or LCZ in N2a cells. 10 µM of EYH or LCZ were added 24 hours after siRNA treatment, and cells were harvested 24 hours after EYH/LCZ treatment. **(G)** 48, 72, and 96 h pretreatment of *Trim66* siRNA at a dose five-fold higher (50 nM) do not ablate the PrP-lowering effects of EYH or LCZ in cells either. 10 µM of EYH or LCZ were added 48, 72, or 96 h after siRNA treatment, and cells were harvested 24 h after EYH/LCZ treatment. (B, C, D) Bars indicate the mean and 95% confidence interval of the mean.

A

| Cell line | PrP expression | Cas9 expression | Compound activity |
|---|---|---|---|
| HEK293 | Yes | Not tested | No |
| A375 | Yes | Yes | No |
| HT29 | Yes | Yes | No |
| U251-MG | Yes | Yes | Minimal |
| U87-MG | Yes | Yes | No |

B

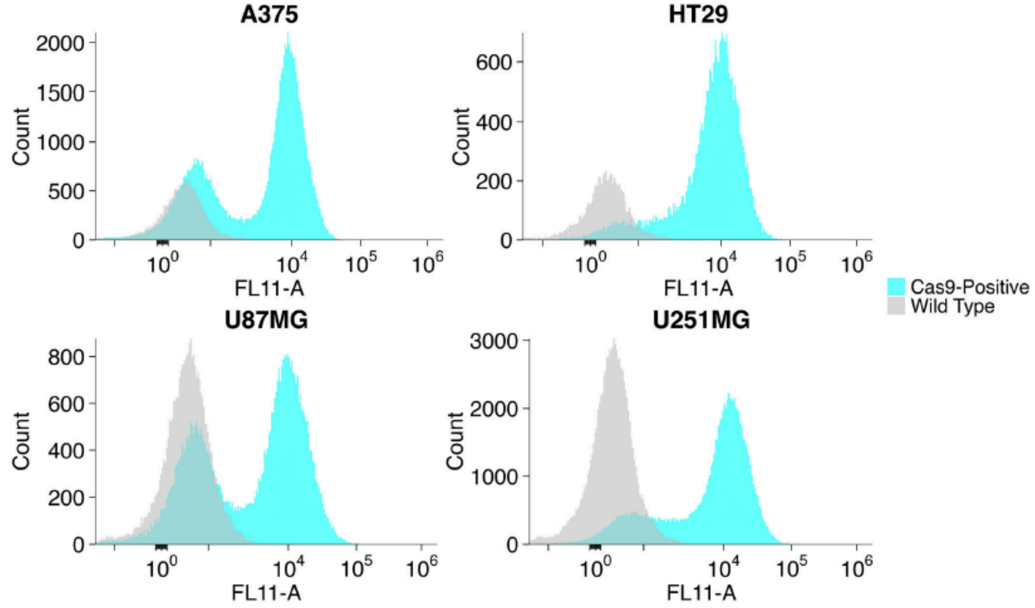


**Figure S3.** Cell lines tested for CRISPR screen compatibility. **(A)** Matrix of human cell lines stably expressing Cas9 summarizing PrP expression, Cas9 expression and compound activity. Note that while all cells express PrP, they are not responsive to the compounds. **(B)** Cas9 activity in A375, HT29, U87MG and U251MG cells analyzed by flow cytometry. Cas9 expressing cell lines were compared to wild-type parental cell lines.

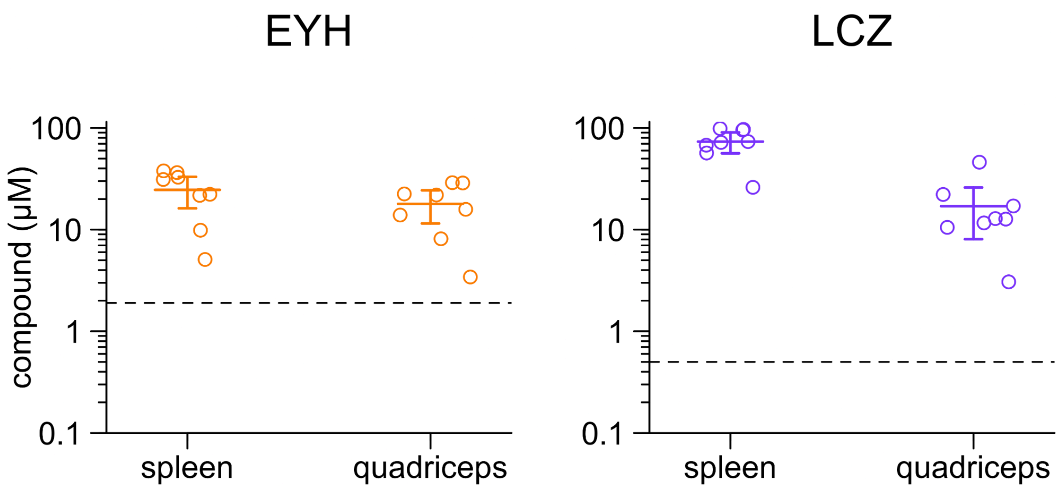


**Figure S4.** *Pharmacokinetics of EYH and LCZ in spleen and quadricep. EYH and LCZ distributed in spleen and quadriceps. Mean concentrations of compounds were 24.6 µM (EYH) and 73.4 µM (LCZ) in spleen and 17.9 µM (EYH) and 17 µM (LCZ). EYH tissue to plasma concentration ratios were 1.5 and 1.0 for spleen and quadricep, respectively. LCZ tissue to plasma concentration ratios were 1.2 and 2.7 for spleen and quadricep, respectively. Dashed lines indicate the $EC_{50}$ of EYH (1.9 µM) and LCZ (0.5 µM) determined in N2a cells.*

**Table S1.** Compounds with corresponding PubChem CID, inchi key, SMILES and CAS numbers.

Supp table compounds.xlsx
